# The phospho-regulated amphiphysin/endophilin interaction is required for synaptic vesicle endocytosis

**DOI:** 10.1101/2023.01.15.524101

**Authors:** Christiana Kontaxi, Michael A. Cousin

## Abstract

The multidomain adaptor protein amphiphysin-1 (Amph1) is an important coordinator of clathrin-mediated endocytosis in non-neuronal cells and synaptic vesicle (SV) endocytosis at central nerve terminals. Amph1 contains a lipid-binding N-BAR (Bin/Amphiphysin/Rvs) domain, central proline-rich (PRD) and clathrin/AP2 (CLAP) domains, and a C-terminal SH3 domain. All domains interact with either lipids or SV endocytosis proteins, with all of these interactions required for SV endocytosis, apart from the Amph1 PRD. In this study, we determined this role and confirmed requirements for established Amph1 interactions in SV endocytosis at typical small central synapses. Domain-specific interactions of Amph1 were validated using *in vitro* GST pull-down assays, with the role of these interactions in SV endocytosis determined in molecular replacement experiments in primary neuronal culture. Using this approach, we confirmed important roles for CLAP and SH3 domain interactions in the control of SV endocytosis. Furthermore, we identified an interaction site for the endocytosis protein endophilin A1 in the Amph1 PRD and revealed a key role for this interaction in SV endocytosis. Finally, we discovered that the phosphorylation status of Amph1-S293 within the PRD dictates the formation of the Amph1-endophilin A1 complex and is essential for efficient SV regeneration. This work therefore identifies an activity-dependent dephosphorylation-dependent interaction that is key for efficient SV endocytosis.

## 1 Introduction

Synaptic transmission results from the activity-dependent fusion of neurotransmitter-containing synaptic vesicles (SVs) at the presynapse, resulting in the activation of cognate postsynaptic receptors. After neurotransmitter release, presynaptic endocytosis is essential to reform SVs, allowing their refilling and recycling for subsequent rounds of neurotransmission. Early studies identified amphiphysin-1 (Amph1), a protein enriched at the presynaptic cytoplasm (Bauerfeind et al. 1997), as a key mediator of SV endocytosis (Shupliakov et al. 1997). This was confirmed when nerve terminals isolated from *Amph1* knockout mice displayed reduced formation of both SVs and endosome-like structures, and fewer competent SVs for repeated rounds of exocytosis (Di Paolo et al. 2002). More recently, immunodepletion of Amph1 *in vivo* resulted in various presynaptic deficits after prolonged stimulation, such as SV shortage, and a reduced number of clathrin-coated vesicles and endosome-like structures (Werner et al. 2016).

Amph1 is a modular protein, with an N-terminal Bin/Amphiphysin/Rvs (N-BAR) domain characterised by the presence of long stretches of amphiphilic α-helices, and can either homodimerize or form heterodimers with another neuronal isoform, Amph2 (Ramjaun et al. 1999; Wigge et al. 1997). The dimeric amphipathic helices of the N-BAR domain recruit Amph1 to highly-curved membranes and exhibit fission-inducing activity (Peter et al. 2004; Snead et al. 2019). The central region of Amph1 contains a clathrin- and adaptor-protein complex 2 (AP2)-binding (CLAP) domain (Slepnev et al. 2000), which facilitates cargo sorting at endocytic pits (Wang et al. 1995) and, when disrupted, inhibits SV endocytosis (Evergren et al. 2004; Jockusch et al. 2005). Finally, the C-terminal Src-homology domain 3 (SH3) of Amph1 interacts with a range of endocytosis proteins that possess a proline-rich domain (PRD) including dynamin-1 (Dyn1) and synaptojanin 1 (Grabs et al. 1997; David et al. 1996; Cestra et al. 1999; Micheva et al. 1997a). Disruption of the Dyn1 interaction inhibits SV endocytosis (Jockusch et al. 2005; Shupliakov et al. 1997), since the Amph1 SH3 domain recruits Dyn1 to endocytosis sites and facilitates fission via regulation of Dyn1 GTPase activity (Takeda et al. 2018).

Amph1 also contains a PRD in its central region, which binds with high affinity to the SH3 domain of endophilin A1 (Micheva et al. 1997b). Similar to Amph1, endophilin A1 is essential for optimal SV endocytosis, since either deletion of its gene or its immunodepletion results in inhibition of the process (Milosevic et al. 2011; Andersson et al. 2010; Ringstad et al. 1999; Gad et al. 2000; Watanabe et al. 2018; Llobet et al. 2011). Endophilin A1 is proposed to have multiple mechanisms during SV endocytosis. For example, its N-BAR domain facilitates recruitment to curved membranes (Andersson et al. 2010) and is associated with membrane remodelling prior to scission (Gallop et al. 2006). Furthermore, it is proposed to regulate both Dyn1-mediated membrane fission (Sundborger et al. 2011; Watanabe et al. 2018; Hohendahl et al. 2017), and uncoating of newly-generated vesicles (Milosevic et al. 2011; Gad et al. 2000). The precise binding sequence within the Amph1 PRD for endophilin has not been defined, however their interaction is inhibited by Amph1 phosphorylation (Murakami et al. 2006; Sekiguchi et al. 2013). Importantly, Amph1 is dephosphorylated during neuronal activity (Bauerfeind et al. 1997; Wigge et al. 1997), with one the major activity-dependent *in vivo* phosphorylation sites (S293) located within the PRD region (Craft et al. 2008), suggesting that this site may control endophilin A1 access to the PRD during SV recycling.

The potential that an activity-dependent Amph1-endophilin A1 complex forms to control SV endocytosis has not been investigated. To address this key question, we performed a detailed molecular mapping of the endophilin A1 interaction site within the Amph1 PRD. We then monitored SV endocytosis in primary cultures of hippocampal neurons previously depleted of endogenous Amph1 and expressing specific full-length interaction-deficient Amph1 mutants. These molecular replacement experiments revealed a key role for Amph1-endophilin A1 interaction in SV endocytosis that is dependent on the phosphorylation status of S293 in the Amph1 PRD.

## 2 Materials and methods

### 2.1 DNA plasmids

Full-length rat Amph1 was obtained from Dr. H. T. McMahon (MRC Laboratory of Molecular Biology, Cambridge, UK). The Amph1-mCerulean (mCer)N1 expression vector was generated by cloning full length rat Amph1 into an mCerN1 vector (generated by replacing EGFP with mCer (Gordon & Cousin 2013)) with the primers 5’-<u>CTCGAGA</u>TGGCCGACATCAAGACGGGCATCT-3’ (sense) and 5’-<u>TGGATCC</u>CGTTCTAGGTGTCGTGTGAAGTTCTC-3’ (antisense) with restriction sites underlined. Point mutations were introduced into Amph1 using standard site-directed mutagenesis protocols with the following primers: 5’-GAGAACATCATCAATT<u>C</u>CT<u>C</u>T<u>AG</u>GGACAACTTTGTACCAG-3’ (sense) and 5’-CTGGTACAAAGTTGTCC<u>CT</u>A<u>G</u>AG<u>G</u>AATTGATGATGTTCTC (antisense) for ^323^FFE^325^ to ^323^SSR^325^, 5’-GGAGACTTTG<u>C</u>TGCATT<u>C</u>G<u>CG</u>CTTTGACCCTTTCAAAC-3’ (sense) and 5’-GTTTGAAAGGGTCAAAG<u>CG</u>C<u>G</u>AAT<u>G</u>CAGCAAAGTCTCC-3’ (antisense) for ^353^DLD^355^ to ^353^HSR^355^, 5’-CAGACATTGCCCT<u>C</u>G<u>CG</u>CTTATGGACGACAAG-3’(sense) and 5’-CTTGTCGTCCATAA<u>GC</u>GCGAGGGCAATGTCTG-3’ (antisense) for ^382^WD^383^ to ^382^SR^383^, 5’-GCCACATACAAA<u>C</u>GCCTCTTTCTAGAGAACTTCACAC-3’ (sense) and 5’-GTGTGAAGTTCTCT<u>A</u>GAAAGAGGC<u>G</u>TTTGTATGTGGC-3’ (antisense) for ^672^GLFP^675^ to ^672^RLFL^675^, 5’-GCATCTCCTGCC<u>G</u>CAGTGCGA<u>G</u>CCAGATCACCTTCAC-3’ (sense) and 5’-GTGAAGGTGATCTGG<u>C</u>TCGCACTG<u>C</u>GGCAGGAGATGC-3’ (antisense) for ^288^PVRP^291^ to ^288^AVRA^291^, 5’-CAAGGAAAGGG<u>G</u>CT<u>G</u>CTGTCGCAGCTCT<u>G</u>CCTAAAG-3’ (sense) 5’-and CTTTAGGCAGAG<u>C</u>TG<u>C</u>GACAG<u>C</u>AG<u>C</u>CCCTTTCCTTG-3’ (antisense) for ^301^PPVPP^305^ to ^301^A_1_A_2_VA_3_A_4_^305^, 5’-GACAAGGAAAGGGGCTGCTGTCCCACCTCTGC-3’ (sense) and 5’-GCAGAGGTGGGACAGCAGCCCCTTTCCTTGTC-3’ (antisense) for ^301^PP^302^ to ^301^A1A2^302^, and 5’-GAAAGGGCCTCCTGTC<u>G</u>CA<u>G</u>CTCTGCCTAAAGTC-3’ (sense) and 5’-GACTTTAGGCAGAG<u>C</u>TGCGACAGGAGGCCCTTTC-3’ (antisense) for ^304^PP^305^ _to_ ^304^A3A4^305^ with modified sites underlined. The Amph1 phosphomutants, S293A and S293E, were generated as previously described (Kontaxi et al. 2022). GST-Amph1 constructs were generated by subcloning Amph1 constructs into a pGEX-KG vector (obtained from Dr. C. Rickman, Heriot-Watt University, Edinburgh, UK) using the primers 5’-CATCAT<u>GAATTC</u>TAGGAGCTCCCAGTGATTCGGGTC-3’ (sense; ΔNBAR), 5’-ATGATG<u>CTCGAG</u>CTATTCTAGGTGTCGTGTGAAGTTCTC-3’ (antisense), and 5’-ATGAT<u>GCTCGAG</u>CTAAGGAGGCAGTTCCTGAGCGG-3’ (antisense; ΔSH3) with restriction sites underlined.

To silence the expression of mouse Amph1, a short-hairpin RNA against the sequence 5’-GGAAGCTTGTGGATTATGA-3’ (shAmph1) was cloned into a modified pSUPER vector (Oligoengine) where the mCer moiety was replaced by synaptotagmin-1-pHluorin (Zhang et al. 2015) using the restriction enzymes BglII and XhoI. A non-targeting control against the sequence 5’-TCGCGATTAGTTCATTAGG-3’ (Scrambled) was generated accordingly. The synaptotagmin-1-pHluorin sequence was replaced in the shAmph1-expressing vector by synaptophysin-pHluorin (sypHy) using the primers 5’-<u>GGATCC</u>ATGGACGTGGTGAATCAGCTGGTG-3’ (sense) and 5’-TG<u>GATATC</u>TTTACATCTGATTGGAGAAGGAGGTG-3’ (antisense) and enzymes BamHI and EcoRV respectively. The sypHy sequence was subsequently mutated to remove a BglII site. All plasmids were verified by Sanger sequencing.

### 2.2 Animals

The wild-type (WT) C57BL/6J mouse colony was utilized for the preparation of primary hippocampal neuronal cultures. Adult female mice were killed by cervical dislocation followed by decapitation, whereas embryos of both sexes were killed by decapitation followed by destruction of the brain at E16-E18. For synaptosomal preparations, adult Sprague Dawley male rats (> 2 months old) were killed by exposure to increasing CO2 concentration followed by cervical dislocation. All experimental procedures were performed according to the UK Animal (Scientific Procedures) Act 1986 on the protection of animals used for scientific purposes and were approved by the Animal Welfare and Ethical Review Body at the University of Edinburgh (Home Office project license to M. Cousin – 7008878 and PP5745138). All animals were maintained on a 12-hour light/dark cycle under constant temperature, with food and water provided *ad libitum.*

### 2.3 Primary neuronal cultures and transfection

Dissected hippocampi were dissociated in papain (10.5 U/ml) and triturated in Dulbecco’s Modified Eagle Medium/Nutrient Mixture F-12 supplemented with 10 % (v/v) foetal bovine serum. Neurons were plated on poly-D-lysine- and laminin-precoated coverslips at 45 x 10^3^ cells/coverslip and maintained in Neurobasal medium supplemented with 0.5 mM L-glutamine, 1 % (v/v) B27 supplement, and penicillin/streptomycin in a humidified incubator at 37 °C/5 % CO2. Cytosine β-D-arabinofuranoside was added at 1 μM on 3 days *in vitro* (DIV) to prevent glial proliferation. Neurons were transfected after 7-10 DIV with Lipofectamine 2000 as per manufacturer’s instructions.

### 2.4 Live-cell imaging and data analysis

Neuronal cultures at 13-17 DIV were mounted in an imaging chamber (RC-21BRFS, Warner) with continuous perfusion of Tyrode’s buffer (119 mM NaCl, 2.5 mM KCl, 2 mM CaCl_2_, 2 mM MgCl_2_, 25 mM HEPES, 30 mM glucose, pH 7.4), supplemented with 10 μM 6-cyano-7-nitroquinoxaline-2,3-dione (CNQX) and 50 μM DL-2-amino-5-phosphonopentanoic acid (AP5). Electrical field stimulation (1-ms pulse width, 100 mA current output) was applied at 10 Hz for 30 s. Live-cell imaging was performed using a Zeiss Axio Observer D1 inverted epifluorescence microscope (Zeiss Ltd., Germany) with a 40x 1.3 NA oil immersion objective. Time-lapse images were acquired at 4 s using a Hamamatsu Orca-ER camera at room temperature. Neurons expressing both sypHy and mCer constructs were visualised at 500 nm, using a 525-nm dichroic and a 535-nm emission filter, respectively. Presynaptic boutons responsive to stimulation were selected and the fluorescence intensity was measured using Fiji is Just ImageJ. Average ΔF/F_0_ calculated for each coverslip was normalised to the maximum fluorescence intensity during stimulation without prior background subtraction. A one-phase exponential fit was used to correct baseline for bleaching that was subtracted from all time points. Distance to the baseline at a fixed time point was used as a measure of the endocytosis extent.

### 2.5 Immunocytochemistry

Neuronal cultures at 14-15 DIV were fixed with 4 % (w/v) paraformaldehyde/PBS and quenched with 50 mM NH_4_Cl/PBS. Following permeabilization with 0.1 % (v/v) Triton X-100 in 1 % (w/v) bovine serum albumin/PBS and blocking in 1 % (w/v) bovine serum albumin/PBS, neurons were incubated with rabbit Amph1 (1:250; Synaptic Systems, cat no. 120 003) and chicken GFP (1:5000; Abcam, cat no. ab13970) primary antibodies for 2 h at room temperature. After washing with PBS, Alexa Fluor secondary antibodies (1:1000; Molecular Probes, Thermo Fisher Scientific) were applied for 2 h at room temperature in the dark. The coverslips were mounted on slides with Mowiol mounting reagent. Neurons were visualised using a Zeiss Axio Observer Z1 inverted epifluorescence microscope (Zeiss Ltd., Germany) and a 40x 1.3 NA oil immersion objective at 475 nm and 555 nm excitation wavelengths. Fluorescent light was detected at 500-550 nm and 570-640 nm using a 495-nm and a 570-nm dichroic filter, respectively. Images were acquired using a Zeiss AxioCam 506 camera and Zeiss ZEN 2 software. Data analysis was performed using Fiji is just ImageJ. Amph1-silenced neurons were visualized at 475 nm excitation wavelength using the anti-GFP antibody as described above. To measure Amph1 expression, regions of interest were drawn around transfected cell bodies (as indicated by GFP staining) and average Amph1 intensity was calculated and normalised to that of untransfected cell bodies. Background was subtracted in all cases.

### 2.6 Crude synaptosome isolation

Adult rat brains were homogenized in ice-cold 0.32 M sucrose, 5 mM EDTA (pH 7.4) after removing the cerebellum. The homogenate was centrifuged twice at 950 x *g* for 10 min at 4 °C and the supernatants were combined. After spinning at 20,400 x *g* for 30 min at 4 °C, the pellet (crude synaptosomal fraction) was collected and lysed in the presence of protease inhibitors (Sigma-Aldrich, cat no. P8849).

### 2.7 Pull-down assay

Expression of glutathione S-transferase (GST)-fused Amph1 constructs in *Escherichia coli* BL21 DE3 cells was induced with 1 mM isopropyl β-D-1-thiogalactopyranoside (Calbiochem, cat no. 420322) at OD_600nm_ 0.6-0.8. After centrifugation at 5000 x *g* for 15 min at 4 °C, the pellets were resuspended in ice-cold buffer containing 10 mM Tris, 150 mM NaCl, 1 mM EDTA, pH 8, protease inhibitors, and 1 mM phenylmethylsulfonyl fluoride. Lysozyme (0.0675 μg/μl, Sigma-Aldrich, cat no. L6876), 4 mM dithiothreitol (Sigma-Aldrich, cat no. D0632), and 10 % (v/v) Triton X-100 were also included. The cells were sonicated at 10 kHz and the clear lysates were spun at 17,420 x *g* for 10 min at 4 °C. The supernatant was transferred to pre-washed Glutathione Sepharose 4B beads (GE Healthcare, cat no. GE17-0756-01) and left rotating overnight at 4 °C. A small volume of the bead-coupled GST-fusion proteins was loaded into a ProbeQuant G-50 Micro Column (GE Healthcare, cat no. 28903408) and washed once in ice cold lysis buffer containing 1 % (v/v) Triton X-100, 25 mM Tris-HCl, 150 mM NaCl, 1 mM EGTA, 1 mM EDTA, pH 7.4, prior to incubation with synaptosomal lysates. The columns were washed successively in ice cold lysis buffer, in NaCl-supplemented lysis buffer (500 mM) and in 20 mM Tris, pH 7.4. Protein elution was achieved by incubating in Laemmli sample buffer for 5 min. All GST-coupled Amph1 constructs were devoid of the N-BAR domain and their total level was estimated with Coomassie Brilliant blue (CBB, Instant Blue Protein Stain; C.B.S. Scientific, cat no. HG73010)) staining prior to immunoblot analysis as previously described (Kontaxi et al. 2022). The primary antibodies used in this study are as follows: goat endophilin A1 (1:1000; Santa Cruz Biotechnology, cat no. sc-10874), goat CHC (1:250: Santa Cruz Biotechnology, cat no. sc-6579), mouse α-AP2 (1:1000; Sigma Aldrich, cat no. A4325), goat Dyn1 (1:500; Santa Cruz Biotechnology, sc-6402), goat syndapin 1 (1:1000; Santa Cruz Biotechnology, sc-10412), and mouse PSD95 (1:1000; BioLegend, cat no. 810401).

### 2.8 Statistical analysis

Statistical analysis was performed using GraphPad Prism 8.4.2 (GraphPad Software Inc). The normality of the data distribution was assessed by performing D’Agostino and Pearson omnibus normality test with significance level set at α = 0.05. Two tailed unpaired *t* test for comparison between two groups or analysis of variance (ANOVA) followed by Tukey’s or Dunnett’s post hoc analysis for multiple comparisons were used for datasets following a Gaussian distribution (presented as mean ± SEM). Mann Whitney test for comparison between two groups or Kruskal-Wallis followed by Dunn’s post hoc analysis for multiple comparisons were used for datasets following a non-Gaussian distribution (presented as median with interquartile range (IQR)). For experiments with a small number of replicates for a normality test to be performed, a parametric test was assumed. Asterisks refer to p-values as follows: *; *p* ≤ 0.05, **; *p* ≤ 0.005, ***; *p* ≤ 0.001, ****; *p* ≤ 0.0001. All experiments consisted of at least three independent biological replicates. Random variation or effect size were not estimated. Sample size and statistical test are indicated in the figure legends.

## 3 Results

### 3.1 Modification of distinct sites on Amph1 eliminates its ability to form complexes with various endocytosis proteins

Amph1 forms interactions with a series of essential endocytosis molecules, such as clathrin, AP2 and Dyn1, which are all essential for optimal SV endocytosis. However, the studies examining SV endocytosis have, to date, been performed in large, atypical synapses (Evergren et al. 2004; Jockusch et al. 2005). To confirm the role of these interactions in SV endocytosis at typical small central synapses, we generated a palette of interaction mutants by first verifying previously identified binding sites on Amph1. To achieve this, mutations were made in Amph1 that was fused to GST to allow extraction of interaction partners from rat brain lysates in pull-down assays. The N-BAR domain was excised from all Amph1 mutants to facilitate their expression in bacteria, since they lack N-BAR proteins in nature (Ren et al. 2006; Bohuszewicz et al. 2016).

First, the binding sites for clathrin heavy chain (CHC) and the α-adaptin subunit of AP2 (α-AP2) within the CLAP domain of Amph1 (Slepnev et al. 2000; Olesen et al. 2008) were modified to generate recombinant GST-tagged Amph1 mutants. The point mutations ^323^FFE^325^→SSR (GST-SSR) were introduced to assess Amph1 binding to α-AP2, whereas the substitutions ^353^DLD^355^→HSR and ^382^WD^383^→SR (GST-HSR/SR) were examined together as the binding site for CHC (Figure 1A). WT and mutant versions of GST-Amph1 baits were then bound to glutathione-sepharose and incubated with rat brain synaptosomal lysates. The interactions of Amph1 with CHC and α-AP2 were then assessed by immunoblotting, in addition to Dyn1 and endophilin A1 (Figure 1B). We confirmed that the Amph1-HSR/SR mutation was sufficient to ablate interactions with CHC (Figure 1C). Similarly, Amph1-SSR lost its ability to bind to α-AP2 subunits (Figure 1D). In addition, the Amph1-HSR/SR had a strong negative impact on α-AP2 binding, whereas Amph1-SSR abolished CHC binding. Importantly, the Amph1-SSR and Amph1-HSR/SR mutants bound both Dyn1 and endophilin A1 to a similar extent as Amph1-WT (Figure 1E, 1F).

**FIGURE 1.**
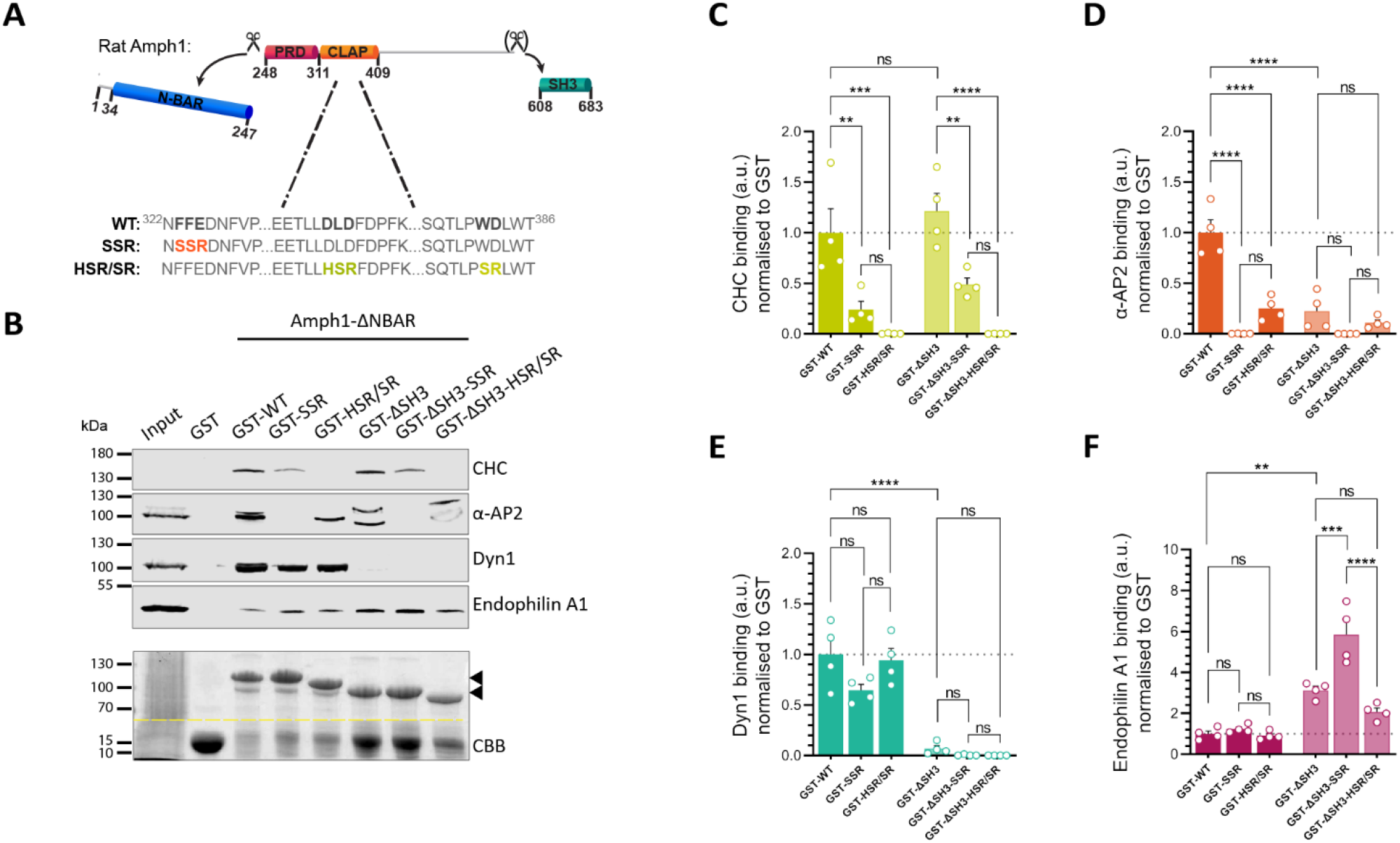
Domain-specific mutations in Amph1 disrupt its ability to form protein-protein complexes. (A) Schematic representation of the structural domains of Amph1 that were used for generating GST-fused N-BAR-lacking mutants either with or without the SH3 domain. Point mutations were introduced within the CLAP domain including FFE to SSR for the α-AP2 binding site; and DLD to HSR and WD to SR for the CHC binding sites as indicated. (B) GST-pull-down experiments were performed using synaptosomal lysates from rat brains and GST-fused Amph1-ΔNBAR mutants. The binding affinity of purified endocytic proteins including endophilin A1, CHC, α-AP2 subunits, and Dyn1 was assessed by immunoblotting. CBB staining was performed in parallel to assess GST-fusion protein levels (arrows). Representative images are displayed. The abnormal separation of α-AP2 subunits may result from the high protein content of the GST-Amph1-ΔSH3 constructs which run at a similar molecular weight. (C-F) Quantification of the binding efficacy of CHC (C), α-AP2 subunits (D), Dyn1 (E), and endophilin A1 (F) by determining relative band intensities normalised to the level of the total GST-fused proteins and WT controls. Bars indicate mean ± SEM. \*\**p* < 0.01, \*\*\**p* < 0.001, and *\*\*\*\*p* < 0.0001 by one-way ANOVA followed by Tukey’s multiple comparison test. *n* = 4 synaptosomal lysates/condition from 4 rat brains.

Amph1 possesses both a PRD and an SH3 domain, which led to the hypothesis that an intramolecular interaction may hinder binding of other ligands to these domains (Farsad et al. 2003). To test this, we generated deletion mutants lacking the SH3 domain. Since Amph1 binds to Dyn1 through this domain (Rosendale et al. 2019; David et al. 1996), it was anticipated that deletion mutants lacking the SH3 domain would be unable to interact with Dyn1. This proved to be the case, with Amph1 devoid of its SH3 domain incapable of binding to Dyn1 (Figure 1E). CHC binding to both WT and CLAP domain mutants was unaffected by the absence of the SH3 domain (Figure 1C), whereas deletion of the SH3 domain severely compromised α-AP2 binding to WT Amph1 (Figure 1D). Finally, removal of the SH3 domain resulted in a significant enhancement of endophilin A1 binding to WT Amph1 and all other CLAP mutants (Figure 1F). Therefore, intramolecular bonds between the Amph1 PRD and SH3 domain may reciprocally control interactions with both α-AP2 and endophilin A1.

### 3.2 CLAP and SH3 interactions are required for Amph1 function in SV endocytosis

The Amph1 mutants characterised above placed us in an ideal position to determine the role of these interactions in SV endocytosis. However, we first confirmed that loss of Amph1 function resulted in defects in this process. To determine this, we generated and validated an shRNA vector against mouse Amph1 (shAmph1) that co-expresses the genetically-encoded reporter sypHy (synaptophysin-pHluorin). SypHy comprises the abundant SV protein synaptophysin with an EGFP that has been engineered to be exquisitely pH-sensitive, inserted into an intralumenal loop (Granseth et al. 2006). Since the interior of SVs is acidic, sypHy fluorescence is quenched in resting neurons, however on SV exocytosis the reporter is exposed to the extracellular environment, resulting in its unquenching. SV endocytosis can be reported by monitoring the requenching of sypHy fluorescence post-stimulation, since the kinetics of endocytosis are rate limiting when compared to SV acidification (Granseth et al. 2006; Atluri & Ryan 2006). A significant knockdown of endogenous Amph1 by shAmph1 was confirmed by immunolabelling in primary hippocampal cultures when compared to scrambled controls (Supplementary figure 1A,1B). When SV regeneration was examined using sypHy, depletion of Amph1 resulted in slower SV endocytosis following stimulation at 10 Hz for 30 s, compared to scrambled controls (Figure 2A,2B). Therefore, loss of Amph1 function results in defects in SV regeneration.

**FIGURE 2.**
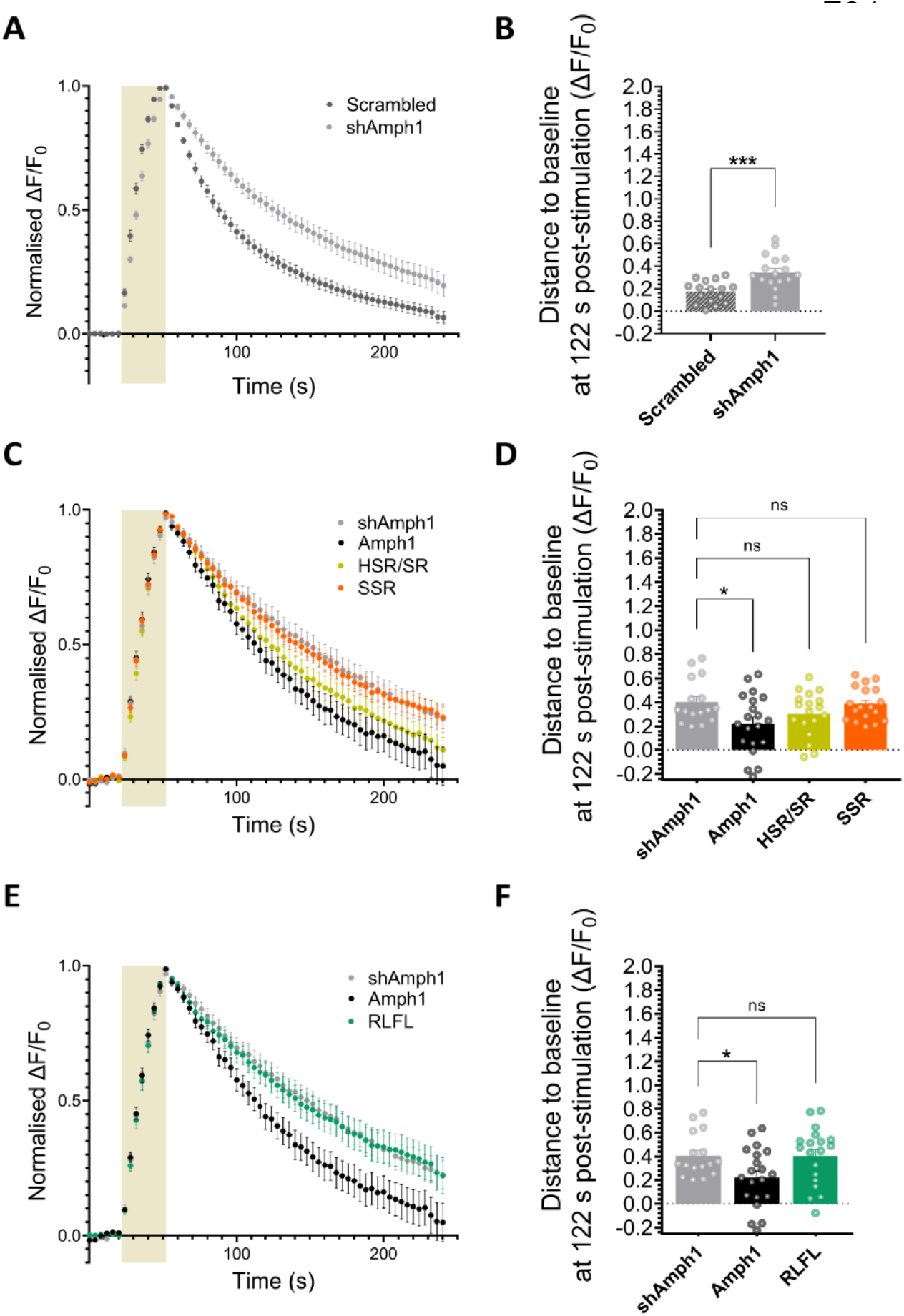
Disruption of Amph1-mediated interactions fails to restore SV endocytosis kinetics in Amph1-depeted neurons. (A,B) Primary hippocampal neurons were transfected with shAmph1 or a scrambled control co-expressing sypHy at 8-10 DIV and imaged at 13-15 DIV. Average sypHy response from neurons stimulated at 10 Hz for 30 s (shaded area) normalised to the stimulation peak (ΔF/F_0_, A) and distance from baseline at 2 min following stimulation (B). Scrambled *n* = 16, shAmph1 *n* =17 coverslips from 4 independent preparations of neuronal cultures, *\*\*\*p* = 0.0007 by unpaired two-tailed *t* test. (C-F) Primary hippocampal neurons from WT mice were co-transfected with shAmph1 (co-expressing sypHy) in addition to mCer (light grey), Amph1 (black), HSR/SR (yellow), SSR (orange), or RLFL (green) mutants at 7-8 DIV and imaged at 13-15 DIV. (C,E) Average sypHy fluorescence response from neurons stimulated with at 10 Hz for 30 s (shaded area) normalised to the stimulation peak (ΔF/F_0_, D,F). Average sypHy fluorescence measuring the distance from baseline at 2 min following stimulation. Bars indicate mean ± SEM. shAmph1 *n* = 16, Amph1 *n* = 20, HSR/SR *n* = 18, SSR *n* = 18, RLFL *n* = 19 coverslips from 5 independent preparations, \**p* < 0.05 by one-way ANOVA followed by Dunnett’s multiple comparison test.

The defect in SV endocytosis observed on Amph1 depletion provides an excellent molecular replacement system to determine which Amph1 interactions are required for optimal SV regeneration. All exogenously-expressed Amph1 rescue constructs for these functional experiments were full-length, to mimic physiological context. Furthermore, mutants were tagged with mCer at their C-termini to allow their visualisation during imaging. We first examined the CLAP missense mutants (SSR and HSR/SR) for their ability to restore SV endocytosis in neurons depleted of endogenous Amph1. All mutants were expressed in the shAmph1-silenced background as confirmed by immunolabelling (Supplementary Figure 2A,2B). Amph1 knockdown neurons expressing both sypHy and Amph1-mCer mutants (or an empty mCer vector control) were subjected to stimulation at 10 Hz for 30 s to evoke SV recycling (Figure 2C). Neurons expressing WT Amph1 had fully restored SV endocytosis kinetics when compared to the empty vector control (Figure 2D). In contrast, neither of the Amph1 CLAP mutants (HSR/SR or SSR) were able to restore SV retrieval kinetics compared to WT (Figure 2D). Therefore, the interactions of either or both CHC and α-AP2 with the CLAP domain of Amph1 are essential for optimal SV regeneration at small central nerve terminals.

The interaction between Amph1 and Dyn1 is essential for SV endocytosis at a number of large, atypical synapses (Jockusch et al. 2005; Shupliakov et al. 1997). We next determined whether this interaction was also required at small central synapses. We avoided truncating the Amph1 SH3 domain to address this question, since we discovered that its removal affects interactions with multiple partners (Figure 1B-F). Instead, we introduced the point mutations ^672^GLFP^675^→RLFL within the SH3 domain that eliminates the interaction with Dyn1 (Grabs et al. 1997). When this Dyn1 interaction mutant was expressed in Amph1-depleted neurons, it failed to restore SV endocytosis kinetics following stimulation at 10 Hz for 30 s (Figure 2E,2F). Therefore, the Amph1-Dyn1 interaction is essential for optimal SV endocytosis at central nerve terminals.

### 3.3 Amph1 interacts with endophilin A1 through a PPVPP motif within its PRD

The Amph1 PRD interacts with the SH3 domain of endophilin A1 (Micheva et al. 1997b) however, the specific interaction site on Amph1 remains unknown. Therefore, to determine the role of the Amph1-endophiln A1 complex in SV recycling, it is essential to pinpoint the exact binding site on Amph1. In contrast to Amph1, the binding site of endophilin A1 on Amph2 has been mapped, and consists of a RKGPPVPPLP motif within its PRD domain (Micheva et al. 1997b). Amongst other roles, Amph1 and Amph2 share common interacting partners including endophilin A1 and, as a result, we hypothesised that Amph1 is likely to associate with endophilin A1 through an analogous motif. Indeed, a similar sequence is present within the PRD of Amph1 (^298^RKGPPVPPLPK^308^). To test the binding affinity of endophilin A1 for this motif, we generated three GST-conjugated versions of Amph1 bearing the point mutations PPVPP→AAVAA (GST-A_1_A_2_VA_3_A_4_), PPV→AAV (GST-A_1_A_2_), and VPP→VAA (GST-A_3_A_4_).

Amph1 is a phosphoprotein with most of its potential phosphorylation sites located within the PRD, including S293, which is its most abundant *in vivo* phosphosite (Craft et al. 2008). We and others have shown that Amph1-S293 undergoes calcineurin-mediated dephosphorylation coupled to neuronal activity, indicating its physiological importance (Craft et al. 2008; Murakami et al. 2006; Kontaxi et al. 2022). Importantly, the phosphorylation status of S293 is critical in controlling the formation of the Amph1-endophilin A1 complex (Sekiguchi et al. 2013; Murakami et al. 2006). For these reasons, we examined two additional GST-fused constructs, a phospho-null and a phospho-mimetic mutant bearing the missense mutations S293A (GST-S293A) and S293E (GST-S293E), respectively. Finally, we mutated another PxxP motif found in the vicinity of this phosphosite for its SH3 recognition potential - ^288^PVRP^291^ →AVRA (GST-AVRA) (Cestra et al. 1999). All constructs were devoid of the N-BAR domain to facilitate bacterial expression, in addition to the SH3 domain to prevent the potential formation of an intramolecular loop (Figure 3A).

**FIGURE 3.**
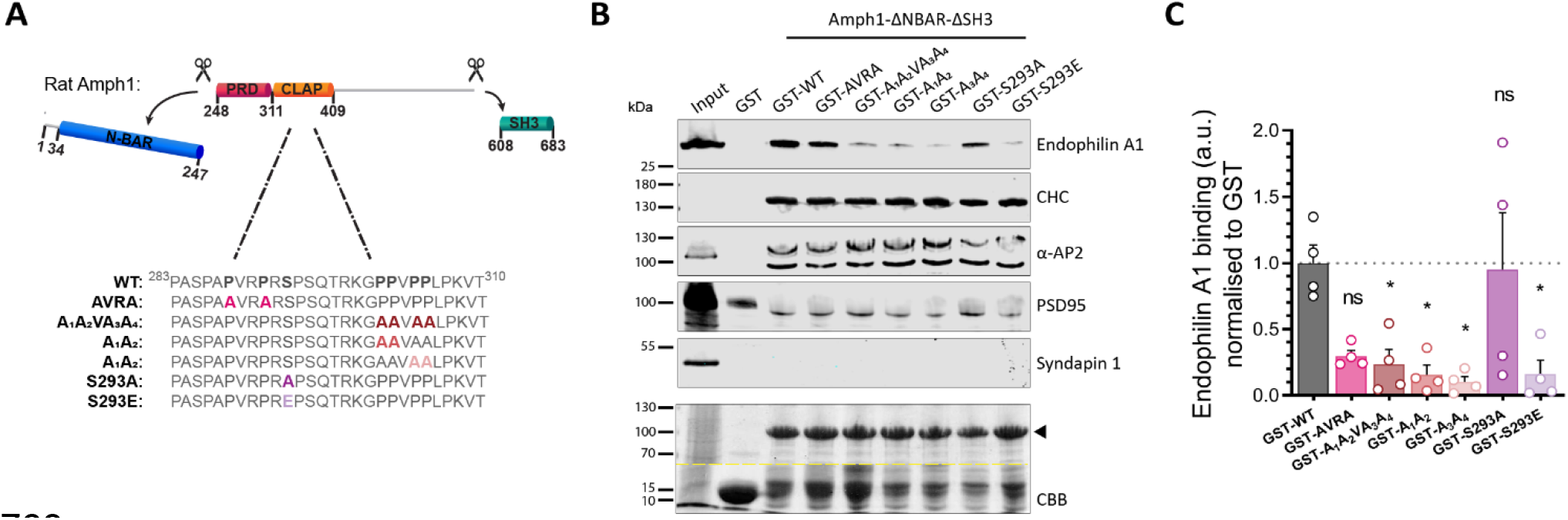
The Amph1-endophilin A1 complex is formed at the motif ^301^PPVPP^305^ within the PRD and is controlled by the phosphorylation status of Amph1-S293. (A) Schematic representation of the structural domains of Amph1 that were used for generating GST-fused ΔN-BAR- and ΔSH3 mutants. Point mutations were introduced within the PRD domain including substitutions of proline to alanine residues for AVRA, A_1_A_2_VA_3_A_4_, A1A2, and A3A4 mutants, and S293A and S293E respectively. (B) GST-pull-down experiments using synaptosomal lysates from rat brains and GST-fused Amph1-ΔNBAR-ΔSH3 mutants to assess binding affinity to endophilin A1 and other SH3-containing proteins, such as CHC, α-AP2, PSD95, and syndapin 1. CBB staining was performed in parallel to assess GST-fusion protein levels (arrows). Representative images are displayed. The abnormal separation of α-AP2 subunits may result from the high protein content of the Amph1-ΔNBAR-ΔSH3 constructs which run at a similar molecular weight. (C) Quantification of the endophilin binding efficacy by determining relative band intensities normalised to the level of the GST-fused proteins and WT Amph1-ΔNBAR-ΔSH3. Bars indicate mean ± SEM. *n* = 4 synaptosomal lysates/condition from 4 rat brains, \**p* < 0.05 by one-way ANOVA followed by Dunnett’s multiple comparison test.

To determine the endophilin A1 interaction site, recombinant GST-tagged Amph1 mutants were used as baits to perform pull-down assays from brain synaptosomal lysates. Their potential interactions with endophilin A1 as well as a number of other SH3-containing molecules, such as PSD-95 and syndapin, were then assessed by immunoblotting (Figure 3B). We revealed that all three PPVPP-related mutants (A_1_A_2_VA_3_A_4_, A1A2, and A3A4) disrupted the binding to endophilin A1 (Figure 3C). The AVRA mutant also exhibits compromised binding, however, this did not reach statistical significance (Figure 3C). These data indicate that endophilin A1 preferentially interacts with the motif ^301^PPVPP^305^ on Amph1 with all the proline residues being variably critical. Importantly, mutation of this site has no impact on either CHC or α-AP2 interactions (Figure 3B).

We next determined the effect of mock phosphorylation of S293 on endophilin A1 binding. We found that the phospho-mimetic mutant S293E abolished the Amph1-endophilin A1 interaction, in contrast to the phospho-null substitution S293A which was not significantly different to GST-Amph1-WT (Figure 3C). As above, these phospho-null and -mimetic substitutions did not impact the association of Amph1 with other SH3-containing endocytosis proteins or CHC and α-AP2 (Figure 3B). Therefore, the phosphorylation status of Amph1 S293 is a key regulator of the Amph1-endophilin A1 complex.

### 3.4 The Amph1-endophilin A1 interaction is required for optimal SV endocytosis

The functional role of the Amph1-endophilin A1 interaction in SV recycling has never been determined. Our generation of Amph1 mutants that selectively ablate endophilin A1 binding allow this question to be addressed for the first time. We therefore examined whether these Amph1 PRD mutants were able to restore defects in SV recycling observed when Amph1 is depleted in central neurons. As before, hippocampal neurons were transfected with shAmph1, or full-length Amph1-mCer with one of the PRD mutants (AVRA, A_1_A_2_VA_3_A_4_, A_1_A_2_, A_3_A_4_) or an empty mCer vector control. All mutants were expressed in the shAmph1-silenced background as confirmed by immunolabelling (Supplementary figure 3A,3B). Neurons were challenged with 300 APs at 10 Hz as before (Figure 4A,4C) and the sypHy response quantified. The AVRA mutant (which did not significantly impact endophilin A1 binding) was able to restore SV endocytosis equivalent to Amph1-WT-mCer (Figure 4A,4B). In contrast, the three other PRD mutants (which did inhibit endophilin A1 binding) could not restore normal SV endocytosis in Amph1-knockdown neurons (Figure 4C,4D). Intriguingly, the A_1_A_2_VA_3_A_4_ mutant, together with A_3_A_4_ mutant, resulted in slightly more pronounced retardation of SV endocytosis, implying a potential dominant-negative role. Taken together, these data reveal that the association of Amph1 with endophilin A1 is required for efficient SV endocytosis at central nerve terminals.

**FIGURE 4.**
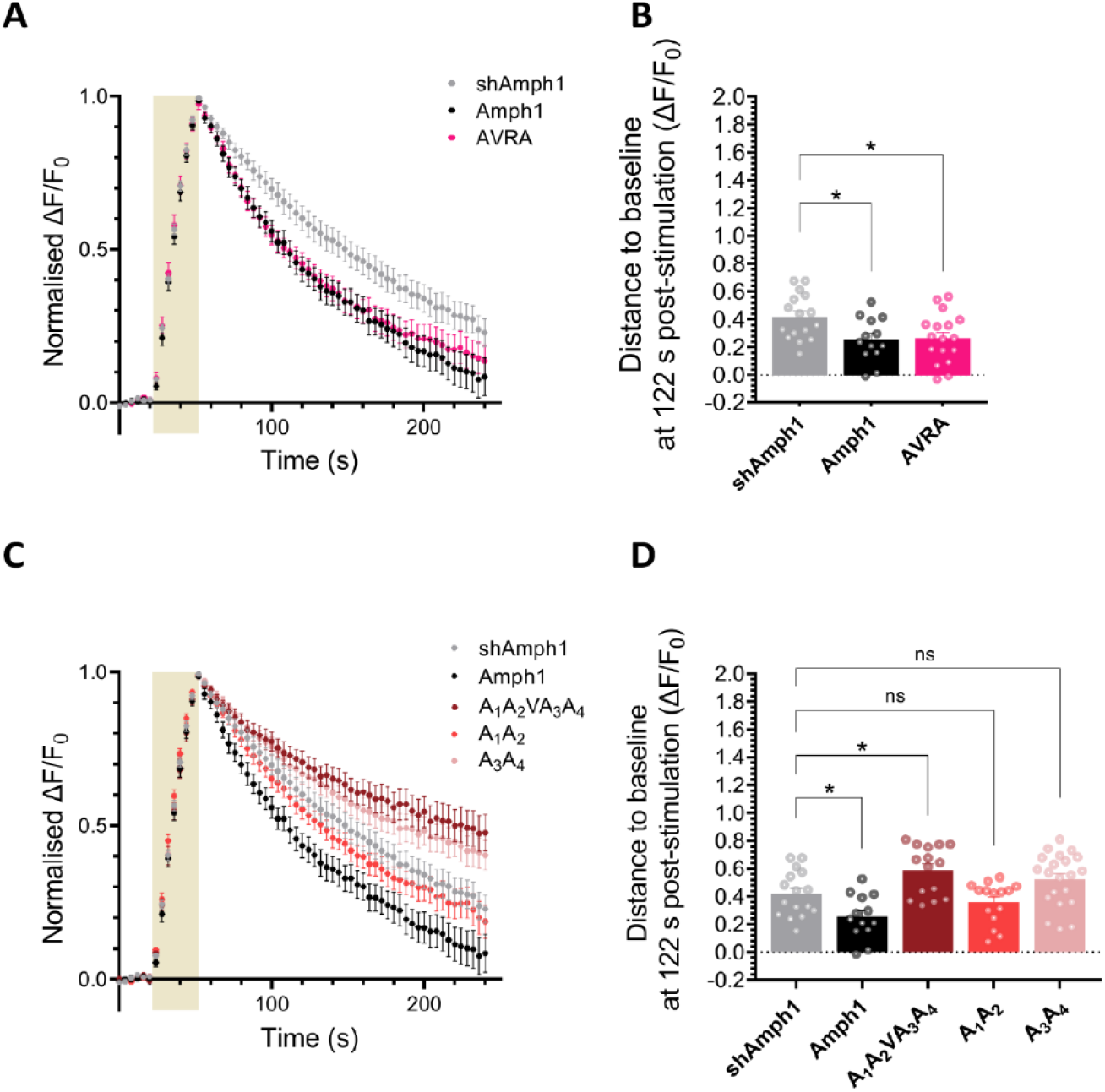
The Amph1-endophilin A1 is essential for efficient SV endocytosis. (A-D) Primary hippocampal neurons from WT mice were co-transfected with shAmph1 (co-expressing sypHy) in addition to mCer (light grey), Amph1 (black), AVRA (fuchsia), A_1_A_2_VA_3_A_4_ (cherry), A1A2 (rose), or A3A4 (salmon) mutants at 7 DIV and imaged at 13-15 DIV. (A,C) Average sypHy fluorescence response from neurons stimulated with at 10 Hz for 30 s (shaded area) normalised to the stimulation peak (ΔF/F_0_). (B,D) Average sypHy fluorescence measuring the distance from baseline at 2 min following stimulation. Bars indicate mean ± SEM. shAmph1 *n* = 16, Amph1 *n* = 13, AVRA *n* = 17, A_1_A_2_VA_3_A_4_ *n* = 15, A1A2 *n* = 15, A3A4 *n* = 21 coverslips from 5 independent preparations, \**p* < 0.05 by one-way ANOVA followed by Dunnett’s multiple comparison test.

### 3.5 The phosphorylation status of S293 is essential for Amph1-mediated SV endocytosis

The Amph1 phospho-mutants, S293A and S293E, differ in their ability to bind to endophilin A1, implying that the activity-dependent phosphorylation status of Amph1-S293 regulates SV recycling through the assembly/disassembly of the Amph1-endophilin A1 complex. Therefore we next determined whether Amph1 S293 phosphorylation controlled SV endocytosis. First, to examine any dominant-negative effect of these phospho-mutants, hippocampal neurons were co-transfected with sypHy and either S293A or S239E to assess their impact on SV endocytosis following stimulation at 10 Hz for 30 s (Figure 5A). In parallel experiments, sypHy was co-expressed with either Amph1-WT-mCer or mCer empty vector (Figure 5A). Overexpression of Amph1-WT-mCer had no dominant-negative effect on SV endocytosis when compared to the empty vector control (Figure 5B). Similarly, overexpression of either S293A or S293E did not alter SV endocytosis kinetics in WT neurons (Figure 5B). Therefore the phospho-null and -mimetic S293 mutants are not dominant-negative.

**FIGURE 5.**
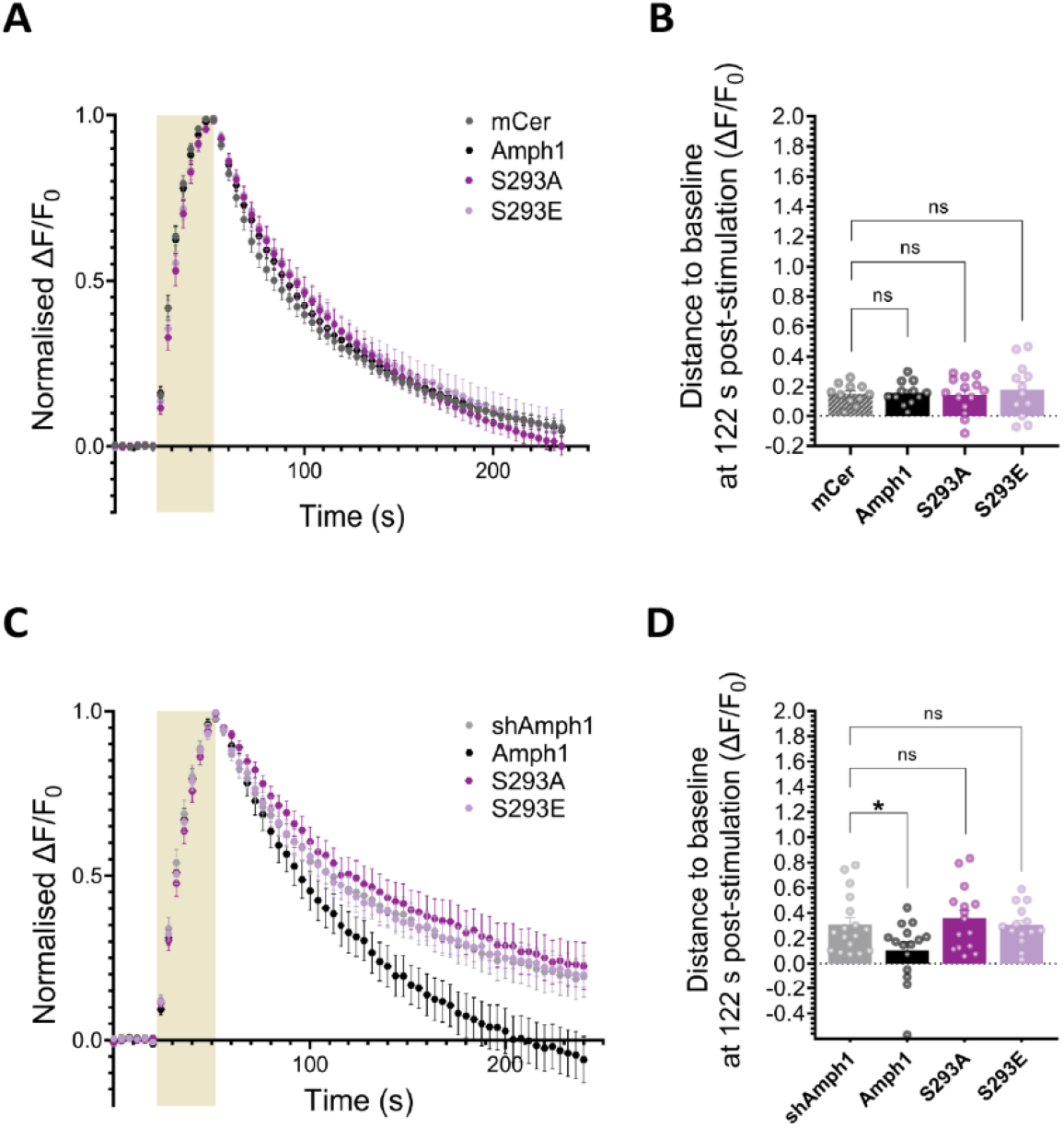
The phosphorylation status of Amph1-S293 controls SV endocytosis. (A,B) Primary hippocampal neurons from WT mice were co-transfected with sypHy and either mCer (dashed grey), Amph1 (black), S293A (purple) or S293E (lilac) at 8-9 DIV and imaged at 13-15 DIV. Average sypHy fluorescence response from neurons stimulated with at 10 Hz for 30 s (shaded area) normalised to the stimulation peak (ΔF/F_0_). (B) Average sypHy fluorescence measuring the distance from baseline at 2 min following stimulation. mCer *n* = 11, Amph1 *n* = 12, S293A *n* = 14, S293E *n* = 12 coverslips from 5 independent preparations, ns by one-way ANOVA followed by Dunnett’s multiple comparison test. (C,D) Primary hippocampal neurons from WT mice were co-transfected with shAmph1 (co-expressing sypHy) in addition to mCer (light grey), Amph1 (black), S293A (purple) or S293E (lilac) at 8-9 DIV and imaged at 13-15 DIV. (C) Average sypHy fluorescence response from neurons stimulated with at 10 Hz for 30 s (shaded area) normalised to the stimulation peak (ΔF/F_0_). (D) Average sypHy fluorescence measuring the distance from baseline at 2 min post-stimulation. Bars indicate mean ± SEM. mCer *n* = 17, Amph1 *n* = 15, S293A *n* = 15, S293E *n* = 14 coverslips from 5 independent preparations, \**p* < 0.05 by one-way ANOVA followed by Dunnett’s multiple comparison test.

Next, we determined whether the S293A or S293E mutants could restore function in neurons depleted of endogenous Amph1. As before, neurons were co-transfected with shAmph1 and mCer-tagged versions of full-length Amph1-mCer or mCer. All mutants were expressed when assessed by immunolabelling in Amph1-silenced neurons (Supplementary figure 3A,3B). Following stimulation with 300APs at 10 Hz (Figure 5C), neither S293A nor S293E was able to restore SV endocytosis in Amph1-silenced neurons in contrast to WT Amph1 (Figure 5D). Overall, these data reveal that the phosphorylation status of Amph1-S293 is critical for controlling both endophilin A1 interactions and SV recycling at central synapses.

## 4 Discussion

Amph1 is established as a key mediator of SV endocytosis at central synapses, via a series of defined interactions with molecules such as CHC, α-AP2, and Dyn1. Using a molecular replacement strategy, we confirmed that these interactions are critical for optimal SV endocytosis at typical small central nerve terminals. Furthermore, we mapped the interaction site for endophilin A1 within the Amph1 PRD and revealed for the first time that this interaction is essential for efficient SV endocytosis. Finally, we determined that the activity-dependent phosphorylation status of S293 within the Amph1 PRD is critical in controlling its interaction with endophilin A1 and SV endocytosis.

These findings were revealed via the integration of interaction assays using Amph1 fragments that were fused to GST as bait, coupled to molecular replacement experiments where full-length Amph1 mutants were expressed in neurons in which endogenous Amph1 was depleted. The extent of knockdown with shAmph1 was variable when quantified via immunofluorescence, however the impact of SV endocytosis was consistent and robust, resulting in defective SV recycling reminiscent of phenotypes observed in Amph1 knockout mice (Di Paolo et al. 2002). This suggests that the levels of endogenous Amph1 present after knockdown are an underestimate and may result from non-specific interactions of the antibody.

### 4.1 CLAP-dependent interactions of Amph1 are essential for efficient SV endocytosis

We confirmed that substitution of the motif ^323^FFE^325^ within the CLAP domain eliminates binding of Amph1 to α-AP2 (Slepnev et al. 2000) but also greatly reduces binding of CHC. Similarly, modification of the sequences ^353^DLD^355^/^382^WD^383^ within the CLAP domain eliminated CHC binding (Slepnev et al. 2000), but also strongly reduced association with α-AP2. These observations reinforce that Amph1-centric complexes are multi-protein, and their formation occurs in a coordinated manner that entails the simultaneous binding of all three partners to ensure the optimal endocytosis (Slepnev et al. 2000).

When examined in our molecular replacement assay, Amph1 mutants that were unable to form complexes with CHC and α-AP2 did not restore SV endocytosis to levels observed with WT Amph1. Therefore, our data support previous observations in large, atypical synapses where CLAP interactions were perturbed via delivery of this domain (Evergren et al. 2004; Jockusch et al. 2005). It is difficult to precisely define the specific roles of CHC and α-AP2 interactions, since binding of both were greatly reduced with both CLAP mutants. However it is likely that both are required, either at the plasma membrane, or more likely, at presynaptic endosomes formed directly by endocytosis. Key roles for both AP2 and clathrin in SV regeneration have been revealed at this compartment over the past decade, especially at physiological temperatures (Watanabe et al. 2014; Soykan et al. 2017; Kononenko et al. 2014; Ivanova et al. 2021; Paksoy et al. 2022). Regardless, our results confirm that CLAP interactions are required for efficient SV regeneration at typical small central nerve terminals.

### 4.2 Amph1 interacts with endophilin A1 to control SV endocytosis

Amph1 and endophilin A1 interact via their respective PRD and SH3 domains (Micheva et al. 1997b; Farsad et al. 2003), however the precise binding site within the Amph1 PRD had not been mapped. We showed that the binding of endophilin A1 to the Amph1 PRD depends on the presence of a class I binding motif within the sequence ^298^**R**KG<u>**P**PV**P**P</u>LPK^308^, which resembles that present on Amph2 (Micheva et al. 1997b). Mutagenesis of the sequence ^301^PPVPP^305^ (underlined above) revealed that both proline pairs, either separately or combined, affect binding of endophilin A1. Exploiting this information, our functional experiments revealed that the first pair ^301^PP^302^ appears to be less critical than the second pair ^304^PP^305^, despite its location at the centre of the class I motif, and in doing so, revealed that this interaction is key for efficient SV endocytosis at central nerve terminals.

In addition, we demonstrated that the phosphorylation status of S293 (which is adjacent to the endophilin A1 interaction motif) is critical for both the control of the endophilin A1 binding and SV endocytosis. Our data support previous observations that the phospho-null mutant S293A associates with endophilin A1, but the phosphomimetic mutant S293E does not (Sekiguchi et al. 2013). Critically, this residue is the prominent *in vivo* phosphorylation site on Amph1 (Craft et al. 2008), and is subject to activity-dependent dephosphorylation by the calcium-dependent protein phosphatase calcineurin (Craft et al. 2008; Murakami et al. 2006; Kontaxi et al. 2022). Therefore, the Amph1-endophilin A1 interaction is coupled to neuronal activity via the action of calcineurin, similar to other key endocytosis molecules that form the dephosphin group of proteins (Anggono et al. 2006; Cousin & Robinson 2001; Lee et al. 2004; Lee et al. 2005).

Both S293A and S293E mutants were unable to restore SV endocytosis in neurons where endogenous Amph1 had been silenced, despite the opposing roles of these phosphomutants in relation to their *in vitro* binding to endophilin A1. It appears that both S293A and S293E are loss-of-function mutants, since their overexpression in a WT background did not impact SV recycling. The loss of S293E function can be explained by its inability to bind endophilin A1 and perform its physiological role. In contrast, the inability of S293A to release endophilin A1 (since it cannot be phosphorylated), may also render this mutant non-functional. If true, this suggests that there is sufficient endophilin A1 present to support endocytosis even if a proportion is sequestered by the S293E mutant. Therefore, the phosphorylation-regulated assembly/disassembly equilibrium of the Amph1-endophilin A1 complex determines SV recycling performance in an activity-dependent manner, most likely via regulation of endophilin A1 clustering at the endocytic pit (Werner et al. 2016).

### 4.3 Intramolecular interactions of Amph1 coordinate SV endocytosis

As discussed, Amph1 shares a number of essential interactions with key endocytosis proteins, including the GTPase Dyn1. In this study, we confirmed that this interaction is mediated via its SH3 domain (Grabs et al. 1997; David et al. 1996) and is essential for SV endocytosis (Shupliakov et al. 1997; Jockusch et al. 2005), most likely via both Dyn1 recruitment (Meinecke et al. 2013; Takei et al. 1999) and Dyn1-mediated fission (Takeda et al. 2018). However, the SH3 domain of Amph1 also appears to perform a key regulatory role via an intramolecular interaction with its PRD (Farsad et al. 2003). Similar intramolecular loops have been reported in other endocytosis proteins, such as endophilin 2 and syndapin 1 (Chen et al. 2003; Rao et al. 2010). For example, binding of Dyn1 to the SH3 domain of syndapin 1 releases an autoinhibition allowing the F-BAR of syndapin-1 to bind membrane (Rao et al. 2010). The increased binding of endophilin A1 in the absence of the Amph1 SH3 domain suggests that Amph1 may be regulated in a similar manner. In this scenario, the attachment of Dyn1 to the Amph1 SH3 domain at the neck of the budding vesicle triggers the recruitment of endophilin A1 to the Amph1 PRD to ensure timely completion of Dyn1-mediated fission (Hohendahl et al. 2017). In support, depletion of Dyn1 results in compromised endophilin recruitment to membranes *in vitro* (Meinecke et al. 2013). Since Dyn1 recruitment depends on the formation of multiple interactions with the Amph1 SH3 domain through distinct motifs (Rosendale et al. 2019), the efficacy of autoinhibitory intramolecular loops to attract Dyn1, especially in an environment that Dyn1 molecules are not stoichiometrically favourable, needs to be assessed further.

Intriguingly, removal of the Amph1 SH3 domain results in compromised interactions with α-AP2 but not CHC. In contrast, Dyn1 is recruited independent to any prior interaction with either α-AP2 or CHC. Therefore, there is potential for the intramolecular interaction between the Amph1 PRD and SH3 domains to generate a binding platform for α-AP2 in addition to previously characterised motifs. The near absence of α-AP2 binding to Amph1 that lacks an SH3 domain suggests that this interaction may only occur when the SH3 domain of Amph1 is unoccupied by external ligands such as Dyn1. This has important implications for the sequence of Amph1-dependent interactions during SV endocytosis with α-AP2 recruitment potentially occurring prior to Dyn1 association, after its dissociation, or both.

In summary, we have revealed a multifaceted role for Amph1 in SV endocytosis at central nerve terminals. We confirmed key roles for interactions via both its CLAP and SH3 domain and revealed that an activity- and phospho-dependent interaction with endophilin A1 is essential for efficient SV endocytosis.

## Acknowledgements

This work was supported by grants from the Loulou Foundation and a College of Medicine and Veterinary Medicine studentship to CK.

## Abbreviations

Amphl: Amphiphysin-1
AP2: adaptor protein complex 2
CBB: Coomassie Brilliant blue
CHC: clathrin heavy chain
CLAP: clathrin and AP2
DIV: days *in vitro*
Dyn1: Dynamin-1
GFP: green fluorescent protein
GST: glutathione S-transferase
N-BAR: N-terminal Bin/Amphiphysin/Rvs
PRD: proline-rich domain
SH3: Src-homology domain 3
SV: synaptic vesicle
sypHy: synaptophysin-pHluorin
WT: wild-type

## 6 Supplementary figures

**SUPPLEMENTARY FIGURE 1.**
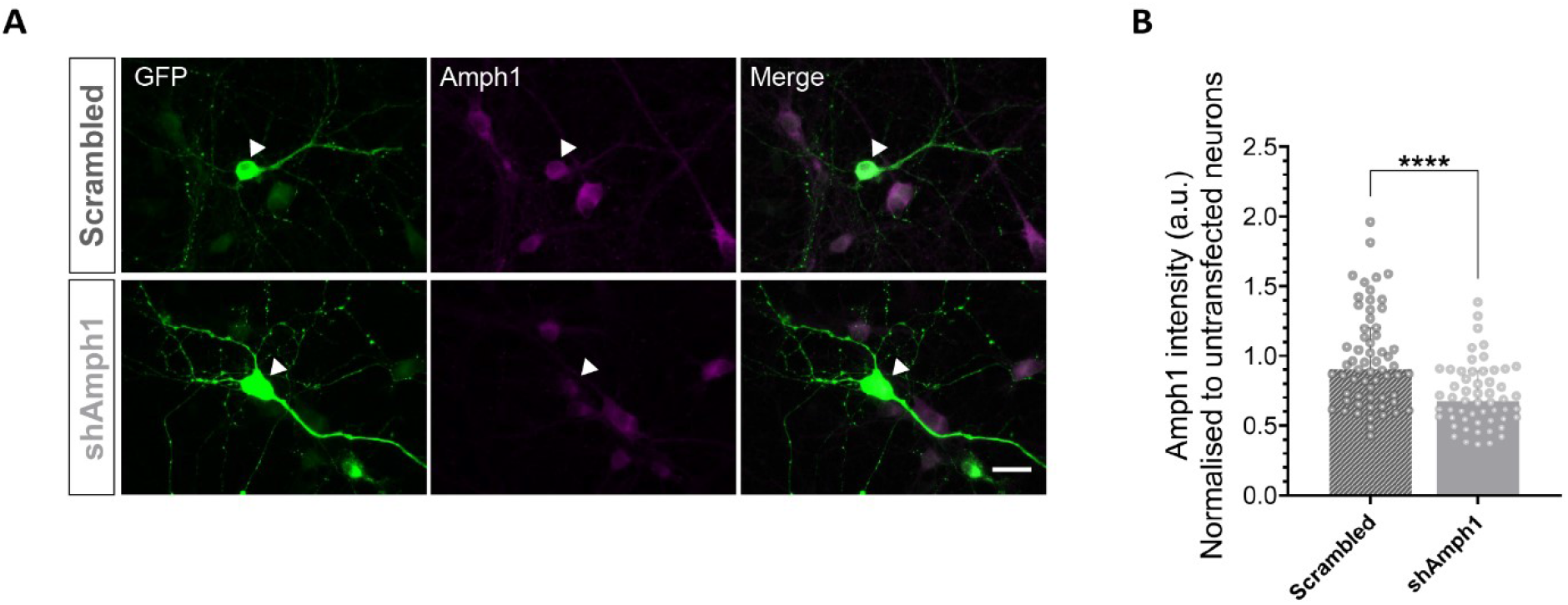
shRNA-mediated silencing of Amph1. Primary hippocampal neurons were transfected with shAmph1 and a scrambled control co-expressing sypHy at 8-10 DIV and fixed at 15 DIV. (A) Representative images of neurons expressing Scrambled and shAmph1 labelled for GFP (green) and Amph1 (magenta). Merged images of GFP and Amph1 are also displayed. Scale bar, 20 μm. (B) Quantification of the mean cell body Amph1 fluorescence intensity (see arrow in A) normalised to untransfected neurons. Background was subtracted in all cases. Bars indicate median with interquartile range. Scrambled *n* = 61 neurons, shAmph1 *n* = 55 neurons from 4 independent preparations of neuronal cultures, \*\*\*\**p* < 0.001 by Mann-Whitney two-tailed test.

**SUPPLEMENTARY FIGURE 2.**
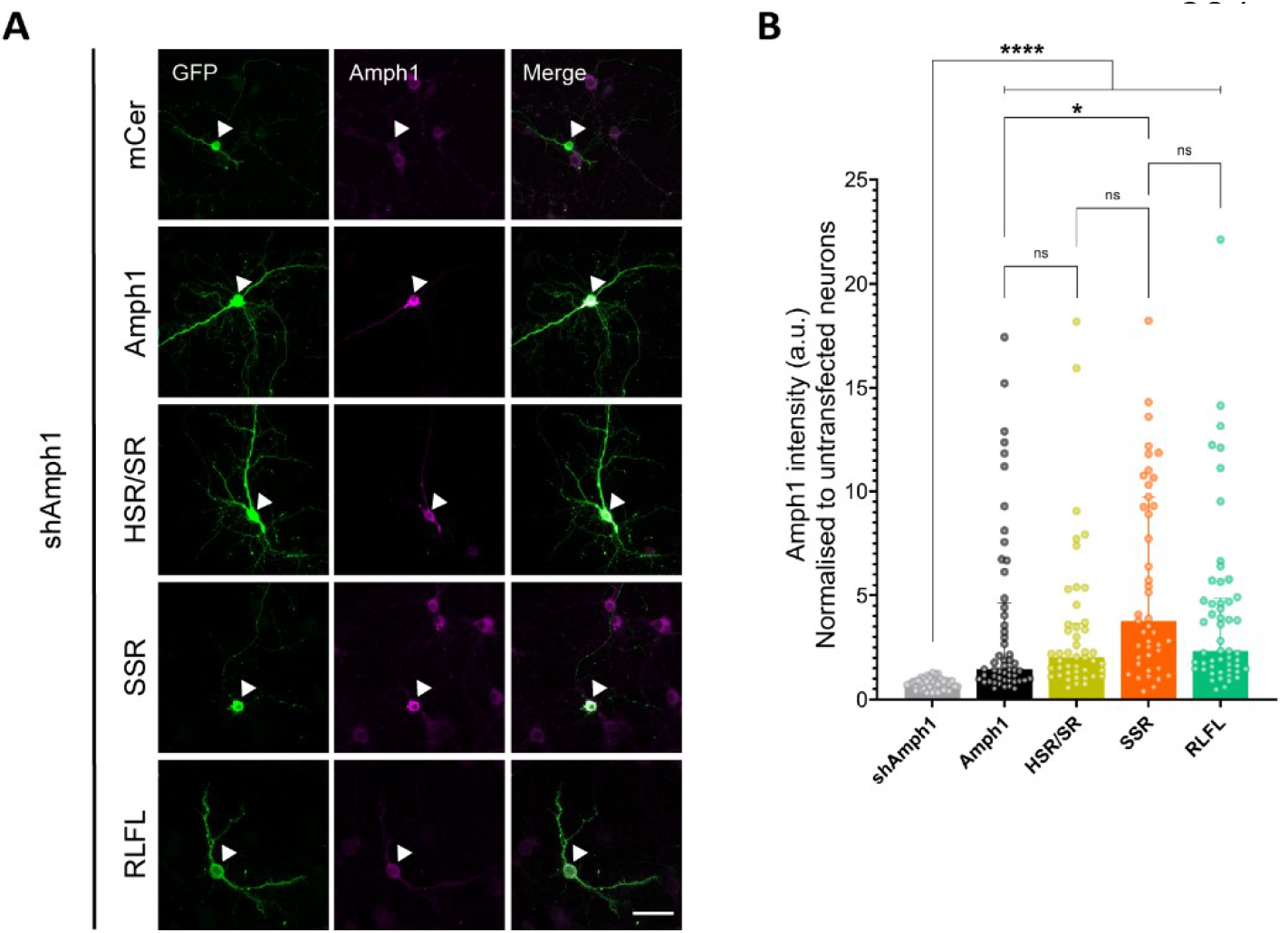
CLAP-localised mutants of Amph1 are expressed in hippocampal cultures. Hippocampal neurons were transfected at 8-10 DIV and fixed at 15 DIV. Representative images of neurons and axons expressing mCer (control) and various Amph1 constructs labelled for GFP (green) and Amph1 (magenta). Merged images of GFP and Amph1 are also displayed. Scale bar, 20 μm. (B) Quantification of Amph1 fluorescence intensity of cell bodies (see arrow in A) normalised to untransfected neurons. Background was subtracted in all cases. Bars indicate median with interquartile range. mCer *n* = 52 neurons, Amph1 *n* = 53 neurons, SR *n* = 45 neurons, SSR/HSR *n* = 43 neurons, RLFL *n* = 52 neurons from 3 independent preparations of neuronal cultures, ns = non-significant, \**p* < 0.05, \*\*\*\**p* < 0.0001 by Kruskal-Wallis test followed by Dunn’s multiple comparison test.

**SUPPLEMENTARY FIGURE 3.**
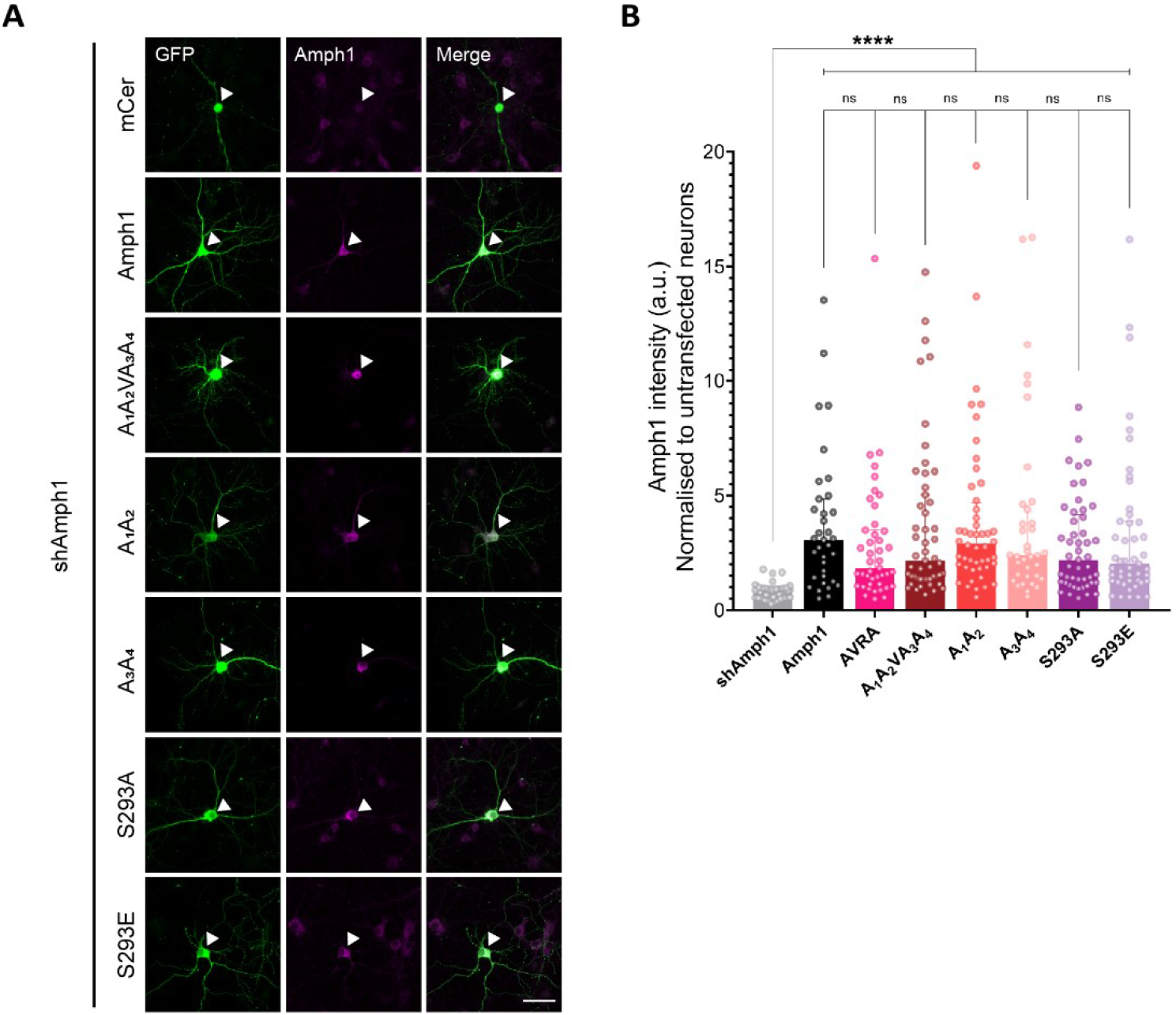
PRD-localised mutants of Amphl are expressed in hippocampal cultures. Hippocampal neurons were transfected at 8-10 DIV and fixed at 15 DIV. Representative images of neurons and axons expressing mCer (control) and various Amph1 constructs labelled for GFP (green) and Amph1 (magenta). Merged images of GFP and Amph1 are also displayed. Scale bar, 20 μm. (B) Quantification of Amph1 fluorescence intensity of cell bodies (see arrow in A) normalised to untransfected neurons. Background was subtracted in all cases. Bars indicate median with interquartile range. mCer *n* = 51 neurons, Amph1 *n* = 35 neurons, AVRA *n* = 43 neurons, A_1_A_2_VA_3_A_4_ *n* = 51 neurons, A1A2 *n* = 48 neurons, A3A4 *n* = 36 neurons, S293A *n* = 49 neurons, S293E *n* = 47 neurons from 3 independent preparations of neuronal cultures, ns = non-significant, *\*\*\*\*p* < 0.0001 by Kruskal-Wallis test followed by Dunn’s multiple comparison test.

